# A novel bio-inspired passive selective cutting mechanism for cutting surgical tools and other material-selective applications

**DOI:** 10.1101/2025.06.10.658779

**Authors:** Martí Verdaguer Mallorquí, Julian Vincent, Andrew Liston, Vladimir Blagoderov, Marc P.Y. Desmulliez

## Abstract

The female sawfly (Insecta: Hymenoptera, Symphyta) uses a double reciprocating saw-like ovipositor to cut into plant tissue, laying its eggs within the cut. We have identified a passive selective cutting mechanism in which the saw discriminates between material properties of the plant tissue without active sensing or external control. The mechanism balances effective cutting with minimal collateral damage to either the plant or the saw. Scaled-up models of the saw cutting into experimental substrates (agar and ballistic gelatine) across a range of stiffnesses reveal an ultimate stress threshold above which substrates are displaced rather than incised. This depends on the interaction between the shape of the saw teeth and the substrate properties and is consistent across multiple species of sawfly. These findings suggest a novel surgical tool design with high selectivity of cutting and reduced collateral damage.

## 1. Introduction

### 1.1 Biomimetics and sawflies

Female ants, bees and wasps (Insecta: Hymenoptera) have an ovipositor through which eggs are laid. This ovipositor has evolved to have other functions, often in parallel with egg-laying, such as stinging (bees and wasps), “drilling” a hole in the substrate (which may be plant or animal material), or cutting with a saw into plant material (sawflies).

The range of substrates that these ovipositors can penetrate, and the inherent simplicity of the mode of action, make them ideal inspiration for biomimetic tools, which has led to multiple studies of very diverse applications ranging from bextra-terrestrial micro-drills for sampling soils to medical devices like self-propelled steerable neurosurgical probes [1,2].

A notable omission from the range of models investigated are the sawflies (Insecta: Hymenoptera, Symphyta) [3]. Insects in this suborder use their saw-like ovipositor (Figure 1) to cut into the soft tissue of plants and lay their eggs inside, without causing excessive damage to the tissues around the cut. Sawfly ovipositors exhibit a high diversity of morphological and structural traits, and are used to oviposit in different parts of various species of plants. It is likely that different ovipositors have evolved to adapt to cutting into specific tissues [4]. A trade-off seems to be at play between the cutting into the plant to lay the eggs, the survival of the plant to enable the larvae to eat the internal plant tissues, and the preservation of the ovipositor from mechanical damage. Learning how ovipositor traits work and relate to the soft tissues they are meant to cut could potentially provide new and effective cutting strategies that are selective and self-preserving. Furthermore, picking the right traits with a proper level of understanding and adapting them for human-made products, could enable the design of cutting tools tailored to the desired substrates.

**Figure 1.**
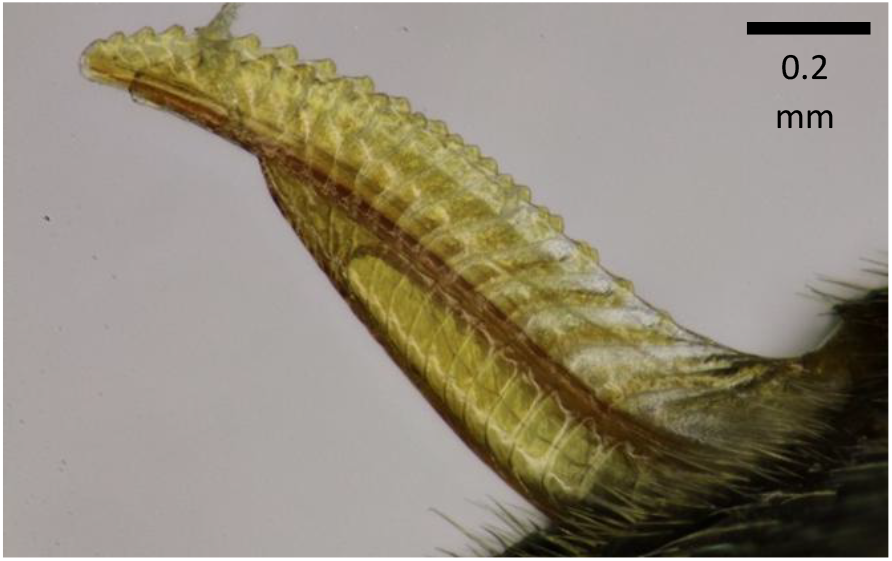
Ovipositor of Sterictiphora geminata (Gmelin, 1790). Adapted from [3]. Open access article (CC-BY 4.0).

As there are over 8000 species in the suborder Symphyta, the amount of knowledge to be gained from an engineering study of these ovipositors is extremely large. In addition, due to the nature of the task performed by the ovipositors, and since biomimetic research on other hymenopterans has yielded knowledge which is highly applicable in designing medical devices, it stands to reason that studying sawfly ovipositors will similarly result in an improvement or innovation of medical devices. The present study addresses this knowledge gap and postulates that some sawflies have adopted a selective cutting mechanism that could be translated for the manufacture of advanced surgical saws and scalpels.

### 1.2 General morphology

The three major parts of the ovipositor are the lance and the two lancets (Figure 2). The vertical plane of symmetry makes the two lancets mirror images of each other.

**Figure 2.**
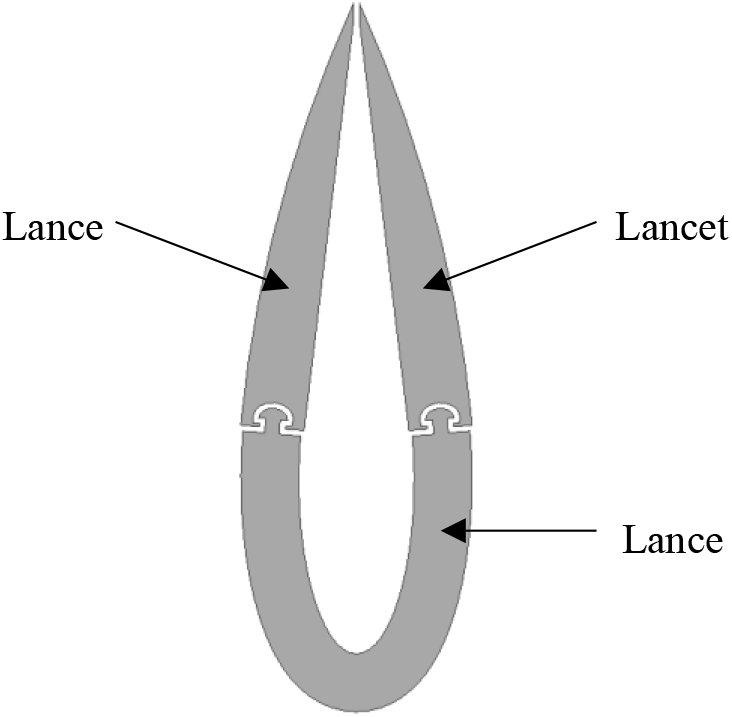
Cross-section of an ovipositor. Adapted from [3]. Open access article (CC-BY 4.0).

During the cutting of the soft tissue of the plant, the two lancets (which bear the teeth) reciprocate while the lance acts as a guide and support for the lancets to slide on (Figure 3).

**Figure 3.**
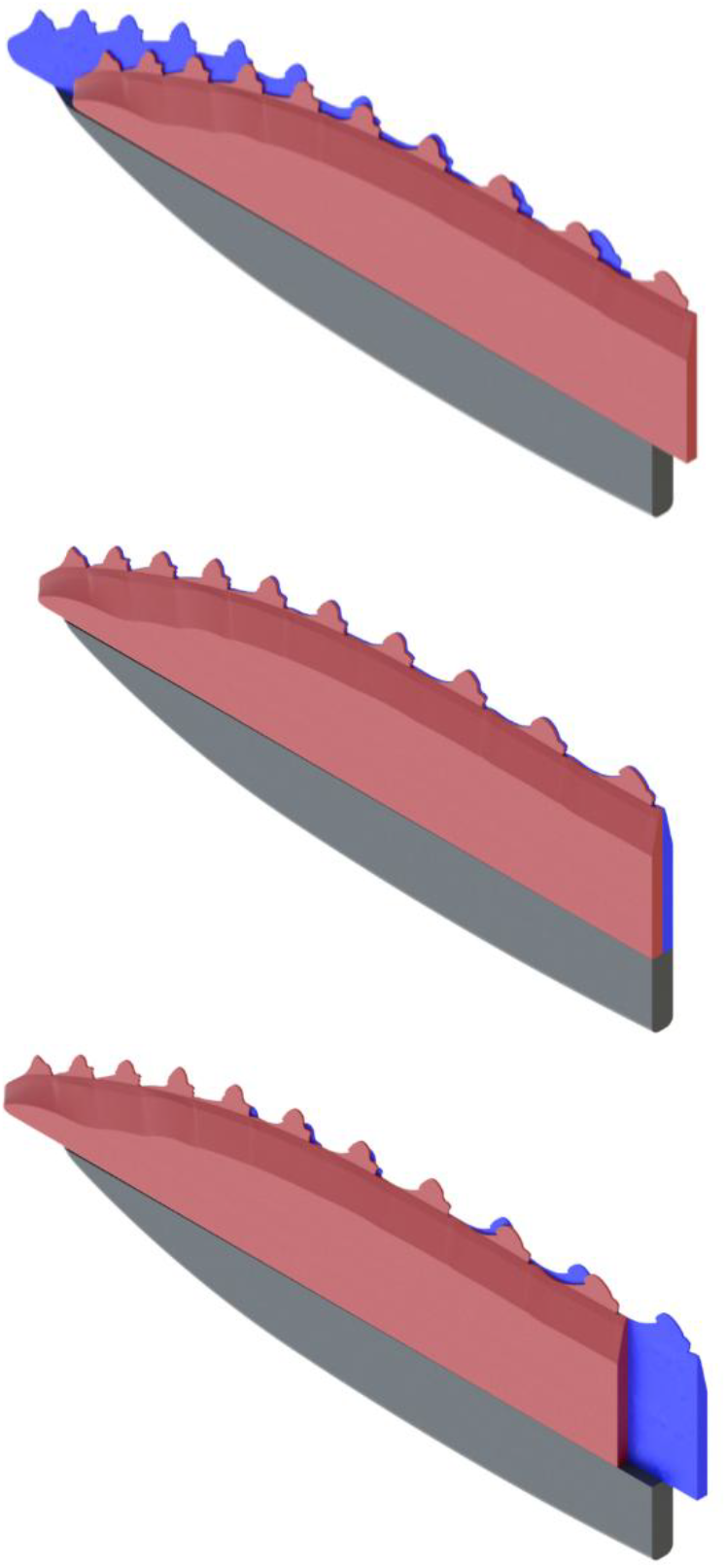
A single cycle of the reciprocating motion.

### 1.2 Concept

The sawfly, *Rhogogaster scalaris* (Klug, 1817), was selected for study as it is readily available, and the plants with which it is associated are known. Later, the ovipositor of the sawfly *Hoplocampa brevis* (Klug, 1816) was also studied, to check that the found principle works across species. The first postulated research hypothesis is that the serrulae and the anterior teeth in this species work together to achieve a synergic effect and enhance the cutting capabilities of the ovipositor. The serrulae (Figure 4) are small teeth on the main saw tooth. The anterior tooth (Figure 4) is a protuberance (the “bump”) at the base of the main tooth [5].

**Figure 4.**
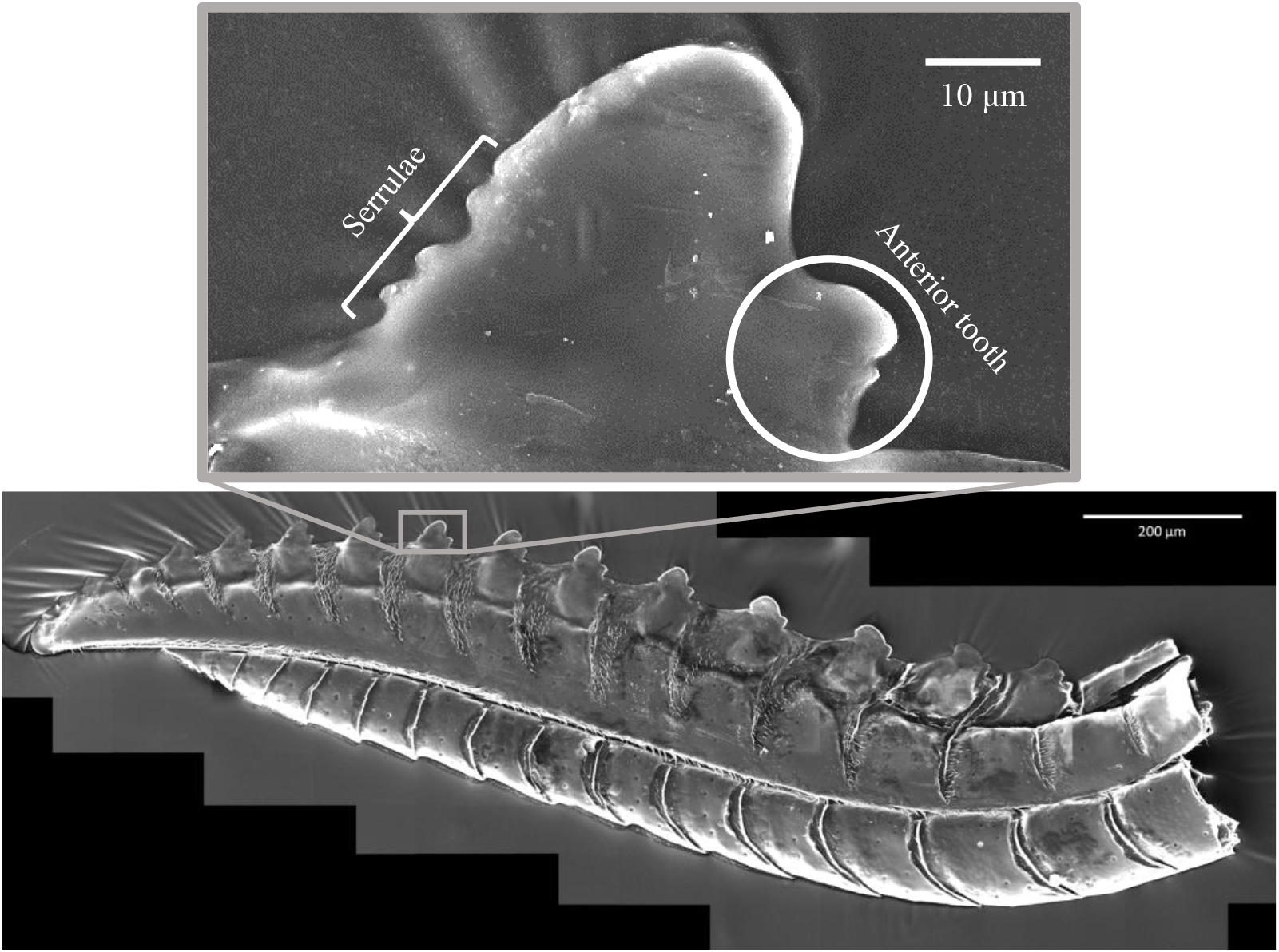
Panoramic and single tooth view of ovipositor. Bottom: panoramic scanning electron microscope (SEM) image of an ovipositor of R. scalaris. Top: a tooth, with serrulae on the apical side and bump on the basal side.

The cutting mechanism is described as follows:

- the two symmetric blades slide against each other in opposite directions,
- the serrulae on a tooth of one blade and the bump on a tooth of the other blade, together trap the substrate, or part of it, and cut it.
- this configuration confines the normal force against the substrate to be cut, and so reduces collateral damage.

The data obtained by differential focus variation profilometry show that the serrulae and the bump are in-plane with each other (Figure 5). This in-plane configuration helped simplify the design of the first test setup.

**Figure 5.**
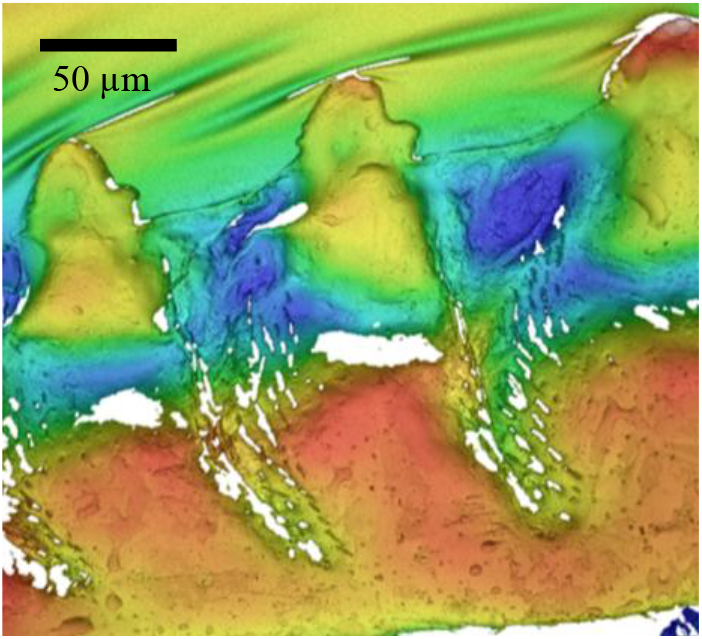
Differential focus variation profilometry of R. scalaris ovipositor.

## 2. Materials and methods

### 2.1 Ovipositors – dissection and examination

The study organisms were *Rhogogaster scalaris* and *Hoplocampa brevis*, two medium-sized sawflies of the family Tenthredinidae, subfamilies Tenthredininae and Nematinae respectively. Specimens, mostly collected in Fennoscandia (*R. scalaris*) and Greece (*H. brevis*), were obtained from the Senckenberg Deutsches Entomologisches Institut. Ovipositors, usually shielded by the third valvula, were dissected under an optical microscope (Signatone S-1160 Probe Station) in 70% ethanol.

To remove residual soft tissue, dissected ovipositors were placed in 15% potassium hydroxide at room temperature for 30 minutes, neutralized in acetic acid and rinsed in distilled water. They were then dehydrated in ethanol using 10% concentration increments (from 70% to 100%), with 30 minutes in each. After overnight immersion in absolute ethanol, they were air-dried and mounted on SEM stubs using conductive adhesive [6] and gold coated using an Agar Scientific sputter coater 108 and a B7390 Gold Target 57 mm x 0.1 mm. The SEM was a TESCAN VEGA operating at 15 kV and a beam current of 100 pA under high vacuum. High-resolution panoramic images were generated by stitching tiled micrographs using the system’s built-in software (Figure 4).

A Sensofar S neox optic profilometer was used to obtain the topography of the ovipositors. Focus variation profilometry reconstructed the surface topography from stacked images across varying heights. Post-processed profiles informed the design of test blades.

The confocal laser scanning microscope (CLSM) imaging was performed on a Leica SP8 system with a HC PL APO 20x/0.75 CS2 air objective (Leica 506517). The excitation was provided by a Supercontinuum White Light Laser; excitation at 405 nm was provided by a Leica SP8 405 nm laser, both using a Leica HyD hybrid detector. The system was operated using the commercial software Leica Application Suite X (LAS X). The carrier slides were 25 mm diameter No. 1.5H high precision glass coverslips (Marienfeld Superior 0117650) held in an Attofluor Cell Chamber (Thermo Fisher A7816). Image stacks (settings in Table 1) were processed into maximum intensity Z-projections using ImageJ, and 3D visualizations were rendered in 3D Slicer. The CLSM samples did not undergo the potassium hydroxide treatment.

**Table 1.**
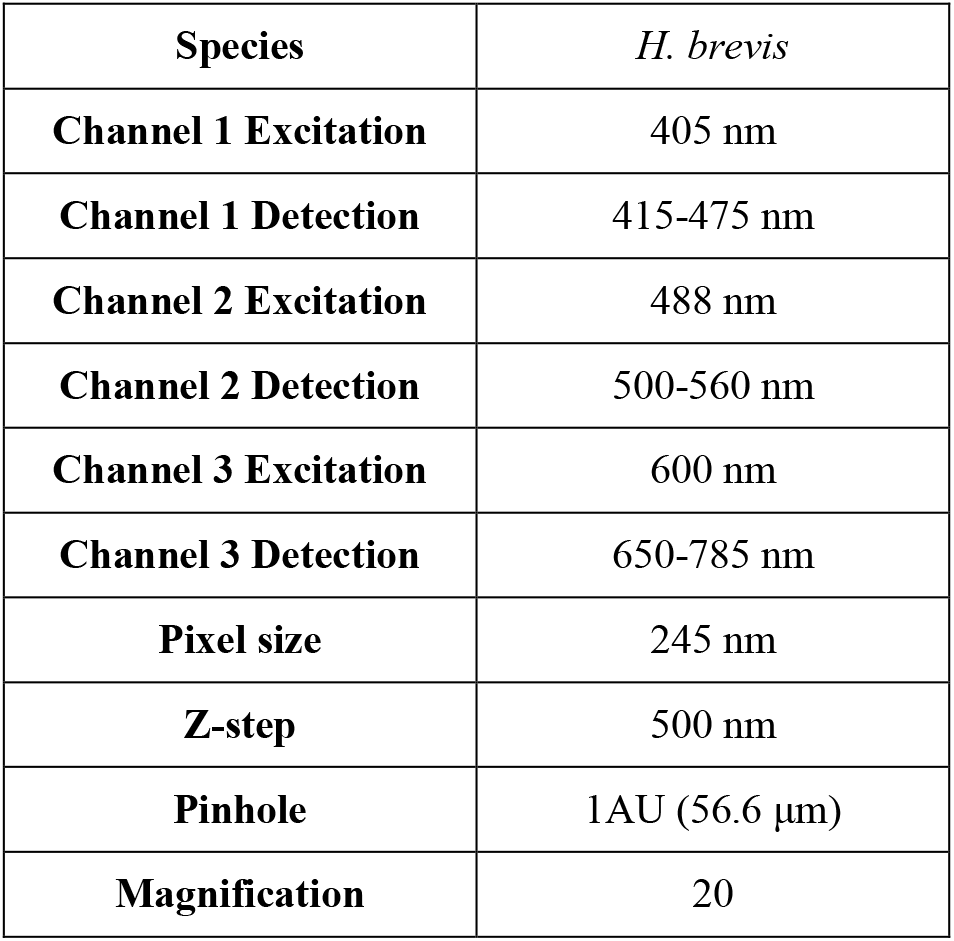
CLSM settings.

### 2.2 Experimental test setup

A representative tooth (9^th^ tooth, counted from the saw apex) from the middle section of a *R. scalaris* ovipositor was traced from SEM images to generate a test blade profile (Figure 6). The region between this tooth and the next more apical one defined the spacing between teeth. A similar process was applied for *H. brevis* using CLSM data.

**Figure 6.**
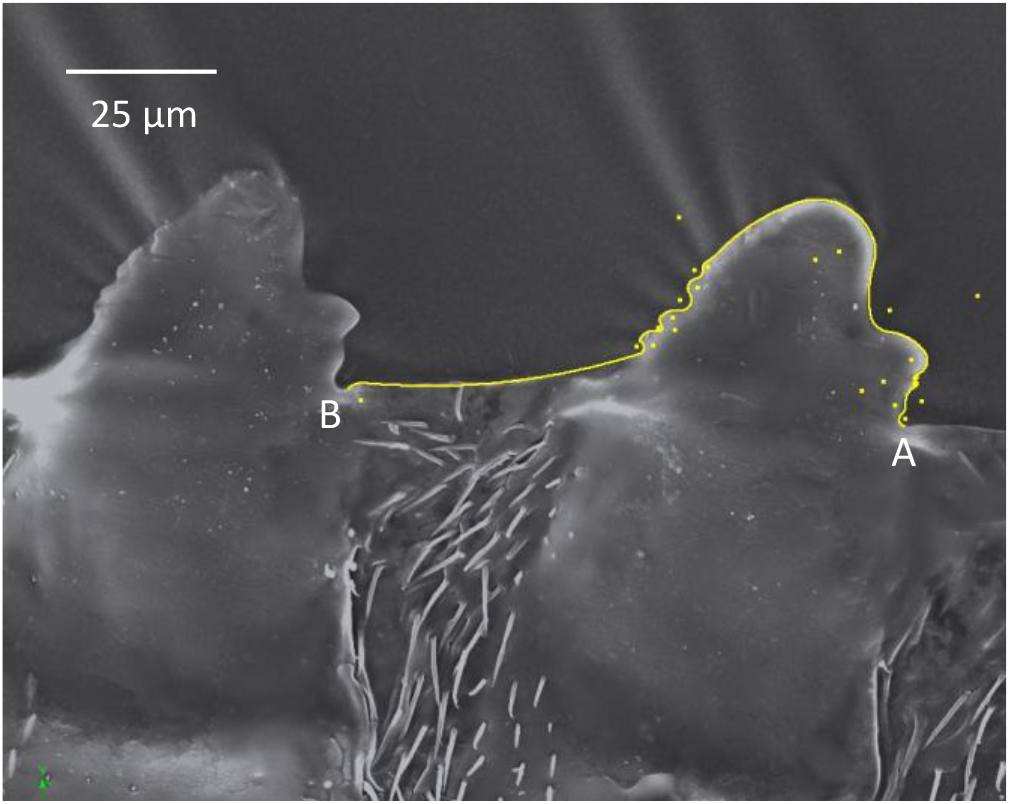
Manually traced profile of the selected tooth (A to B).

An experimental rig was designed and constructed to study the cutting characteristics (Figure 7). A Hounsfield h20k-w traction test machine pulled the blades along the Y-axis, while the substrate rested between the teeth, thereby allowing the control of the degrees of freedom.

**Figure 7.**
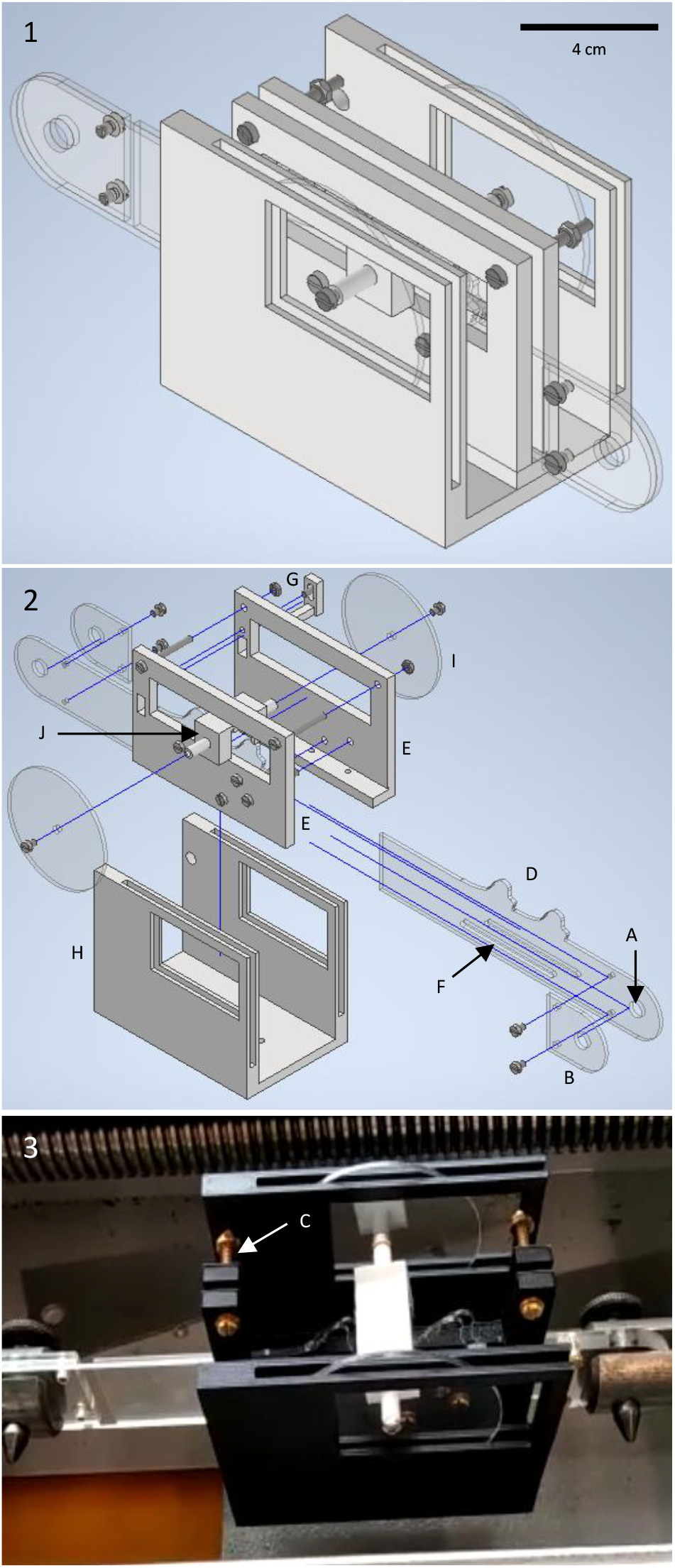
CAD design and produced test rig attached to the traction test machine. Top CAD images show the “closed” set up. The photo shows the “open” set up. The substrate carrier (Figure 8) is white and facing downwards, ready to be cut. A: Attachment hole for securing the blades to the Hounsfield h20k-w test machine. B: Extra material for traction alignment. C: Springs providing constant force to clamp the assembly. D: Blade. E: Core walls enclosing the blades. F: Slotted holes allowing one blade to slide during operation. G: Small arm preventing vertical movement and increasing stability. H: Outer walls of the U-shaped part holding the disc. I: Disc supporting and guiding the substrate, limiting off-plane motion. J: Substrate carrier, connected through wall windows to the internal disc.

The test had two conditions: one which forced the substrate to stay within the range of the teeth (closed) and one which allowed the substrate to move out of the range of the teeth (open).

The blades were mounted to the Hounsfield h20k-w traction test machine via a hole at their extremity and aligned for straight traction using a reinforced section. A spring-based clamping system held the blades between two parallel walls, one blade fixed and one slidable along slotted tracks. This allowed movement while maintaining contact. An upper retaining arm restricted vertical motion and stabilised the assembly. The teeth were visible through a central window. The core assembly was fixed to a U-shaped frame that supported a pair of internal discs with large cut-outs. The substrate carrier was attached to these discs, preventing out-of-plane motion and ensuring that cutting was confined within the intended plane.

The substrate carrier (Figure 8) had openings for the blades to cut the substrate and cylindrical protrusions which allow them to be bolted to the discs (Figure 7).

**Figure 8.**
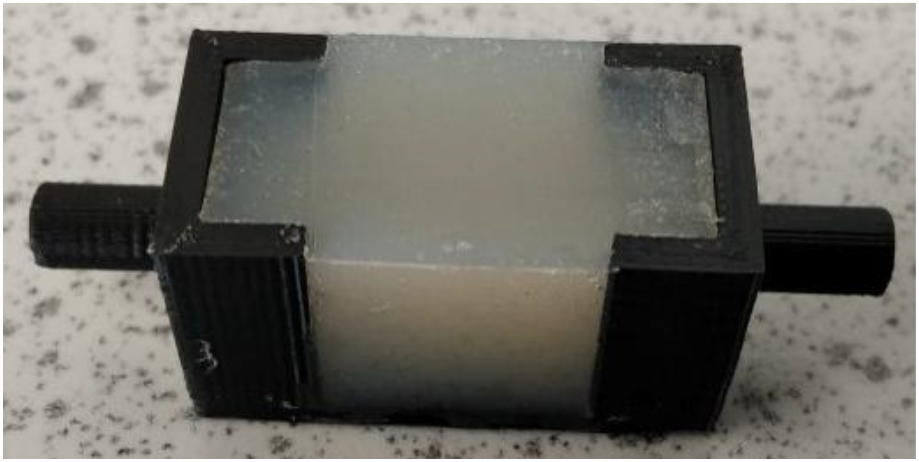
An agar filled substrate carrier. 30 × 15 × 16 mm excluding the cylindrical protrusions, 2 mm walls.

Another version of the test rig allowed direct observation of cutting. The front guide window was removed, and two LEDs provided a light source from behind the substrate (Figure 9).

**Figure 9.**
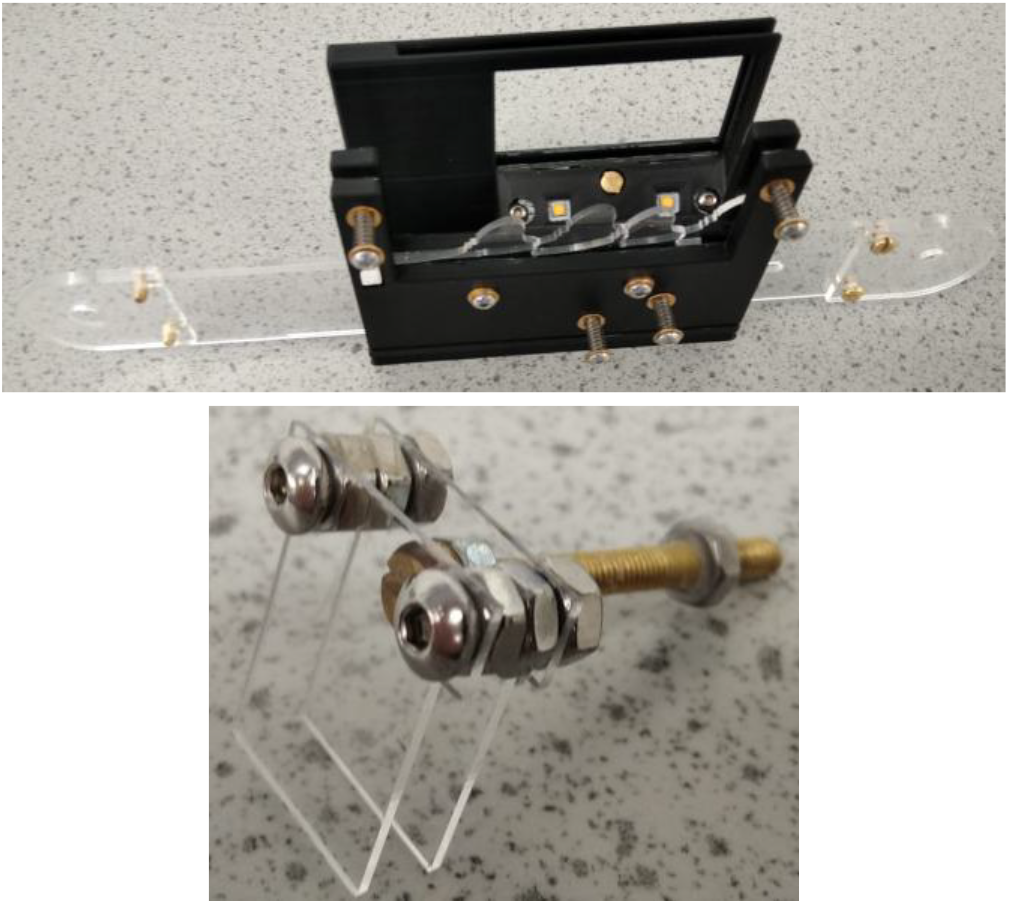
Top: modified test setup allowing direct visualisation of the cutting process. Bottom: new container design for thin substrate slices.

The container for the substrate was replaced by two PMMA transparent covers sufficiently separated for a thin slice of substrate to be formed between them (Figure 9) and cutting was recorded with a digital camera and macro lens.

### 2.3 Substrate preparation

Substrates of agar and ballistic gelatine were prepared for the cut and compression tests (Table 2). Agar powder with water was heated to boiling in a microwave oven, then cooled down at room temperature. Ballistic gelatine was prepared using porcine skin gelatine powder (type A, 300 Bloom) at a concentration of 10% w/w, in accordance with the Federal Bureau of Investigation (FBI) ballistic gelatine standards [7].

**Table 2.**
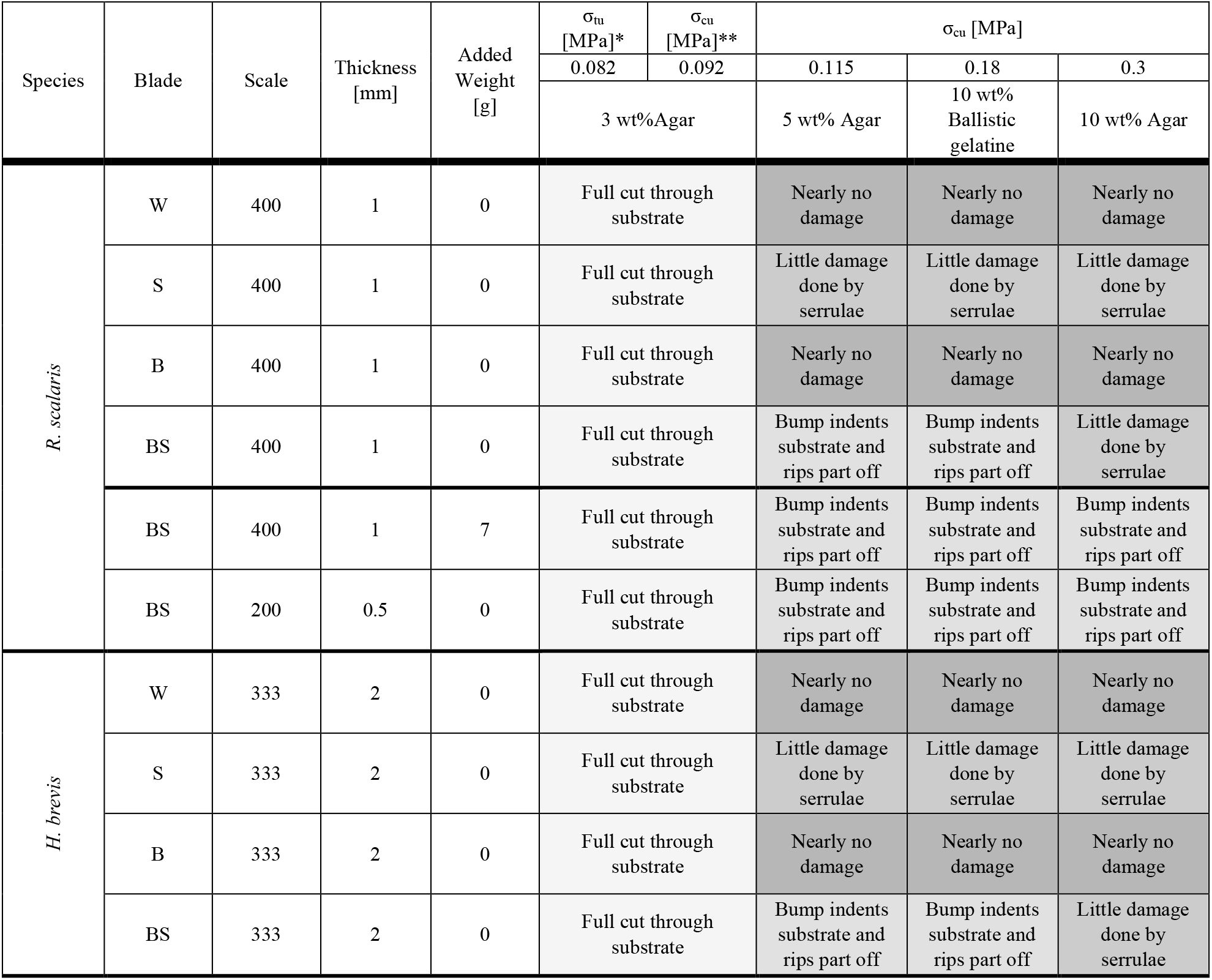
Cutting test results across all conditions, including agar and ballistic gelatine concentrations and their corresponding ultimate compressive stresses, blade configuration tests results, added weight and scale change tests results, and different species comparision. σ_tu_= ultimate tensile stress, σ_cu_= ultimate compressive stress. *Value from literature [9], **Extrapolated value from acquired data.

The mixtures were then poured into moulds designed for three tests: cut tests, 2D cut tests, and compressive tests. To ensure consistency, substrates for a test series were cast using the same batch of material. The cut test mould consisted of PMMA slabs enclosing the substrate carrier. The 2D cut test used a PLA container with a through-hole matching the carrier profile, vented for uniform casting. Compression samples were cast in a four-sample PLA mould; dimensions were 22 × 15 × 15 mm. Flatness and demoulding were facilitated using a thin PMMA cover and a light wax coating respectively. All compressive tests were performed using an Instron 5800 universal test machine. For cut tests, the force exerted on the substrate was estimated by weighing the loaded substrate carrier on a laboratory scale.

### 2.4 Manufacturing techniques

All the components of the experimental setups, including mould and substrate carriers, were designed using Autodesk Inventor. All the 3D printed parts were produced in PLA using the Ultimaker S5 filament printer equipped with a 0.4 mm nozzle. An Epilog mini laser plotter was used to cut the PMMA parts.

## 3. Substrates and qualitative tests

### 3.1 Substrates and characterisation tests

The mechanical properties of agar and ballistic gelatine were adjusted by modifying the amount of agar powder or gelatine powder added to the water (Table 2). Ballistic gelatine was selected because of its mechanical resemblance to human soft tissue [7] [8].

To characterise the mechanical properties of the substrates, uniaxial compression tests were performed on samples cast using the moulds described in section 2.3. The samples measured 22 × 15 × 15 mm, and the crosshead speed was 1 mm/min. Six tests were performed for each substrate. The 3% agar substrate was not characterised, but data were obtained that fit extrapolations from the data obtained for other percentages of agar [9].

### 3.2 Cutting tests

Four sets of blades were produced to test the hypothesis that the bump and serrulae work synergistically. The blade profiles (Figure 10) were BS (with bump and serrulae), B (only bump), S (only serrulae) and W (neither bump nor serrulae). The teeth on the blades were isometrically scaled up 400 times, resulting in teeth approximately 13.8 mm tall and 1 mm thick. Each set of blades was tested against the four substrates (3%, 5% and 10% agar, as well as 10% ballistic gelatine). All cut tests were done at a speed of 10 mm/min and repeated four times for each combination of blade and substrate. The substrate carriers, including the substrate, weighed approximately 15 g.

**Figure 10.**
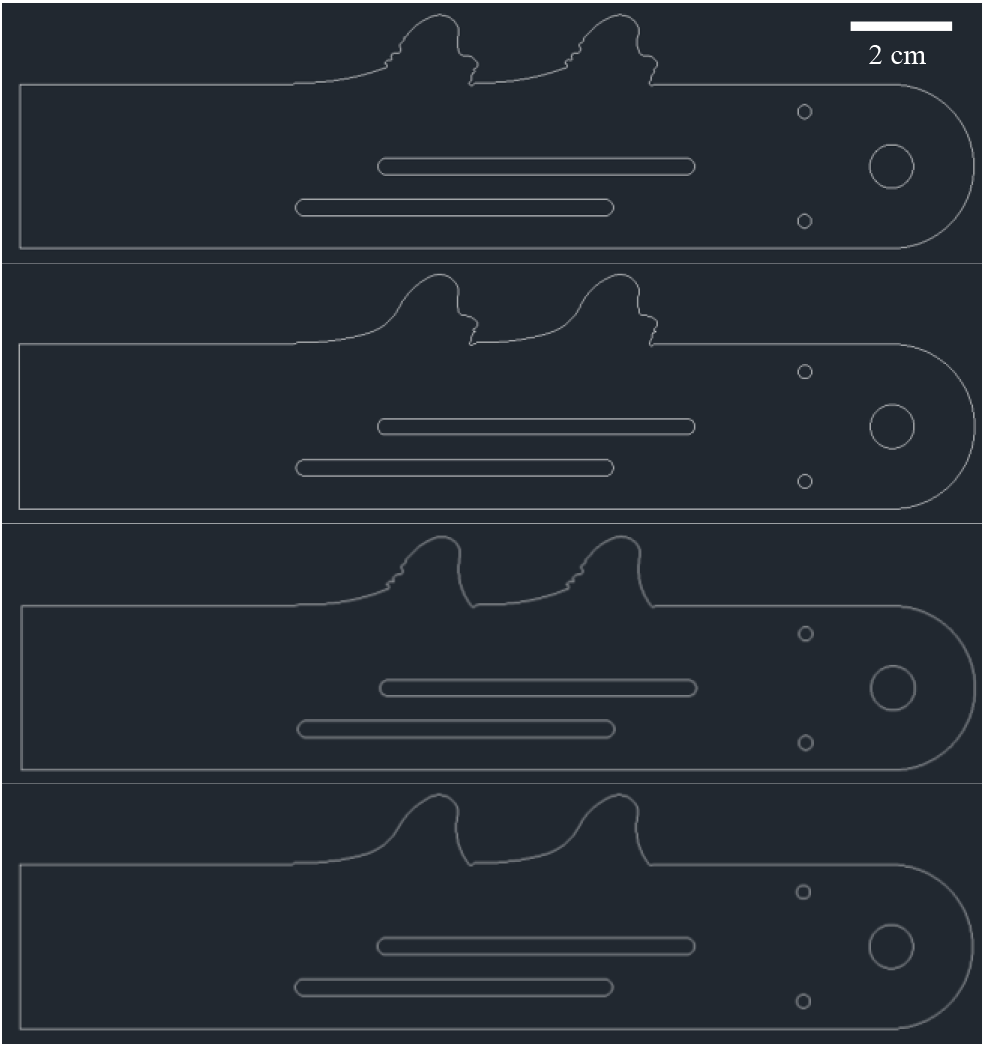
The different teeth shapes used in the tests. From top to bottom: Bump-Serrulae (BS), Bump (B), Serrulae (S), Without both (W).

### 3.3 Added weight and scale change cut tests

The substrates which were not cut would be pushed upwards out of the range of the teeth. There were two main factors: the movement of the substrate and its mechanical failure. To isolate the effects of the force against the substrate and the blade scale, two additional tests were performed: one with weight added to the substrate carrier to increase vertical force, and another with smaller blades to increase stress concentration. Of course, those two are intrinsically related, but this is nonetheless a simple approach to isolating them in the experiments. First the BS set was tested against all substrates with weight added to the substrate carrier to increase the force pushing the substrate against the blades. The total substrate package weight in this case was approximately 22 g. Another set of BS blades at 200 times the size of the model tooth had teeth about 6.9 mm tall and a thickness of 0.5 mm. This new set of blades was also tested against all the agar and ballistic gelatine compositions.

### 3.4 Different species

A second species of sawfly, *Hoplocampa brevis*, was selected to test that the hypothesis is applicable to other species possessing the necessary traits. In *H. brevis*, the tip of the tooth is considered to be the bump, which should maintain the functionality of the principle involved. The tooth for this set of blades was manually traced from the Confocal Laser Scanning Microscopy (CLSM) image shown in Figure 11.

**Figure 11.**
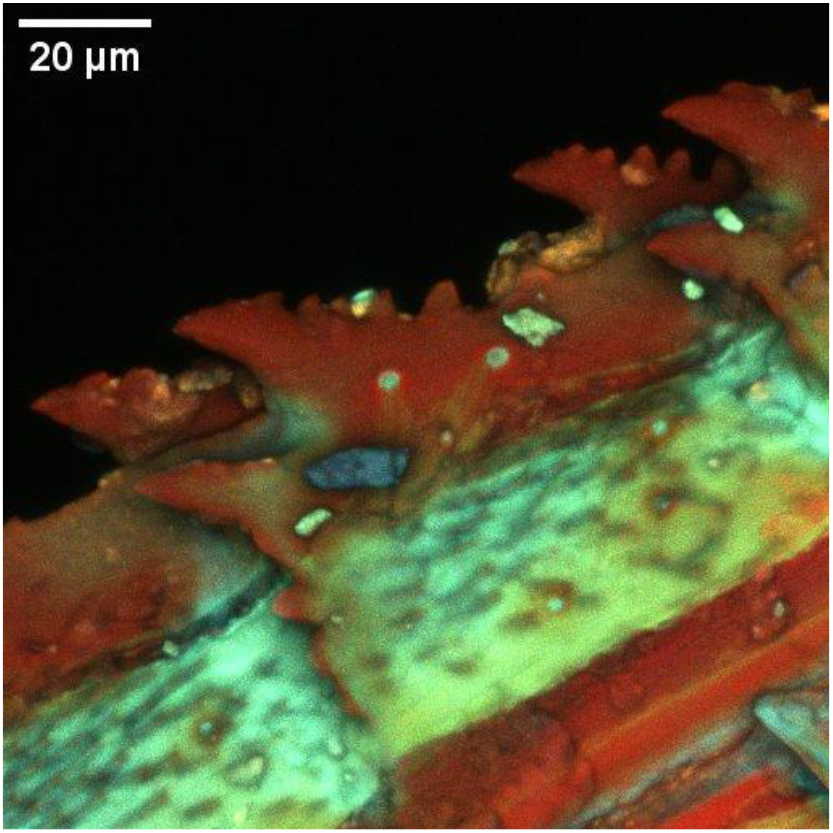
Confocal laser scanning microscopy of H. brevis.

Four sets of blades were produced at an isometric scale of 333.3, yielding a blade thickness of 2 mm and teeth about 6.64 mm tall. The four configurations were akin to those used for *R. scalaris*, (Figure 12). These four sets of blades were tested against the four substrates.

**Figure 12.**
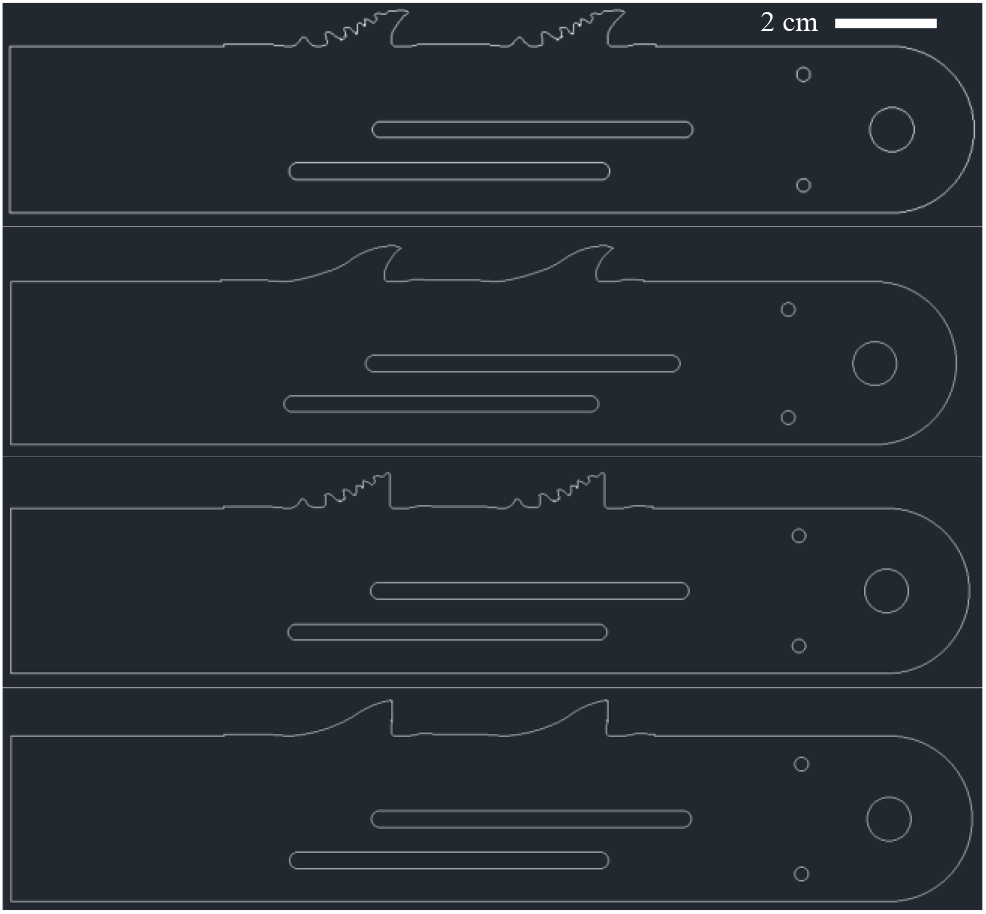
H. brevis sets of blades. Top to bottom: BS, B, S, W.

## 4. Results

### 4.1 Substrates and characterisation tests

Compressive stress-strain curves were obtained for six samples of 5% agar, 10% agar and 10% ballistic gelatine (Figure 13-Figure 15). An averaged curve and ultimate compressive stress were calculated for each material; the latter is summarized in Table 2. The agar curves are approximately linear, while the ballistic gelatine presents a characteristic hyper-elastic behaviour. These results were used to interpret the cut tests.

**Figure 13.**
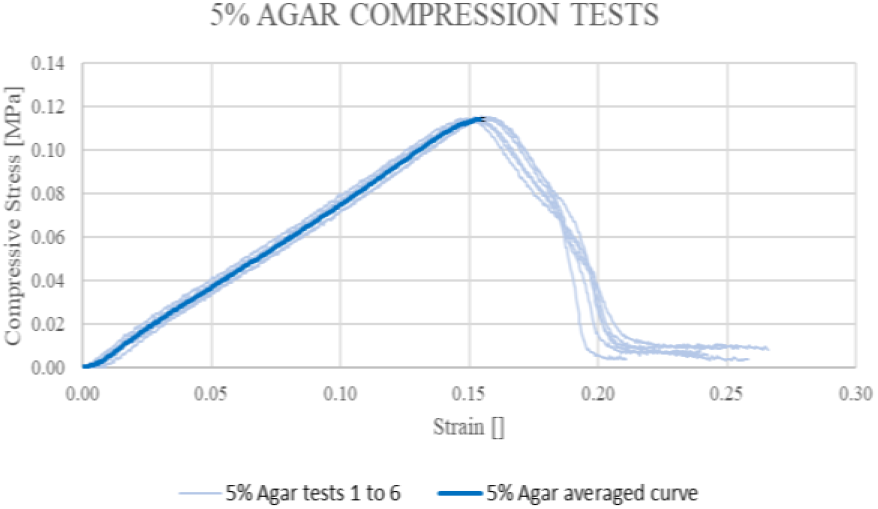
Compressive stress - strain curves for 5% agar.

**Figure 14.**
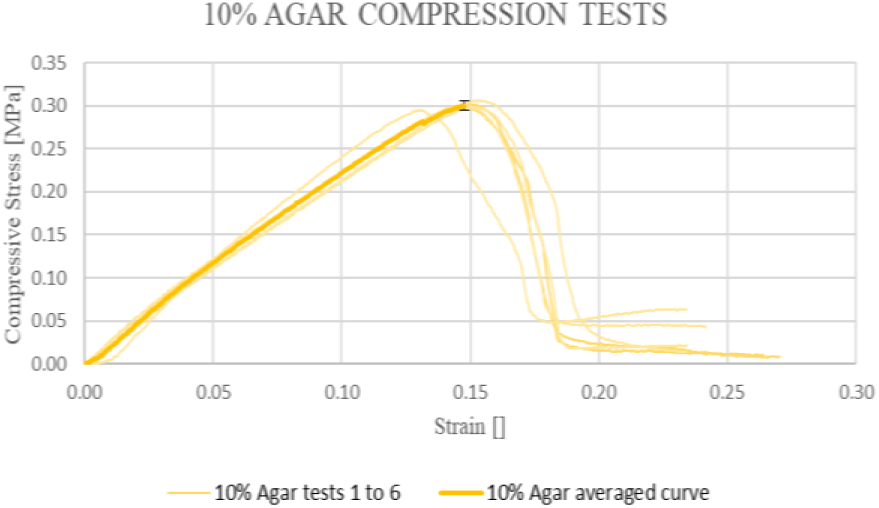
Compressive stress - strain curves for 10% agar.

**Figure 15.**
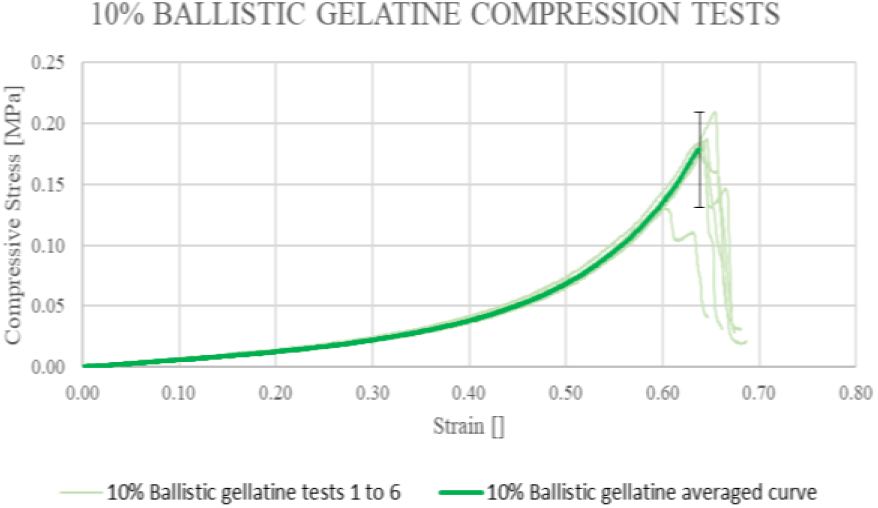
Compressive stress - strain curves for 10% ballistic gelatine.

### 4.2 Cutting tests

The cutting tests using the “open” set up resulted in four types of damage: “full cut through substrate”, “bump indents substrate and rips part off”, “little damage done by serrulae” and “nearly no damage” (Table 2).

All four models of blades produced full cuts through the 3% agar substrates (Figure 16, Figure 17) with minimal differences between blade sets.

**Figure 16.**
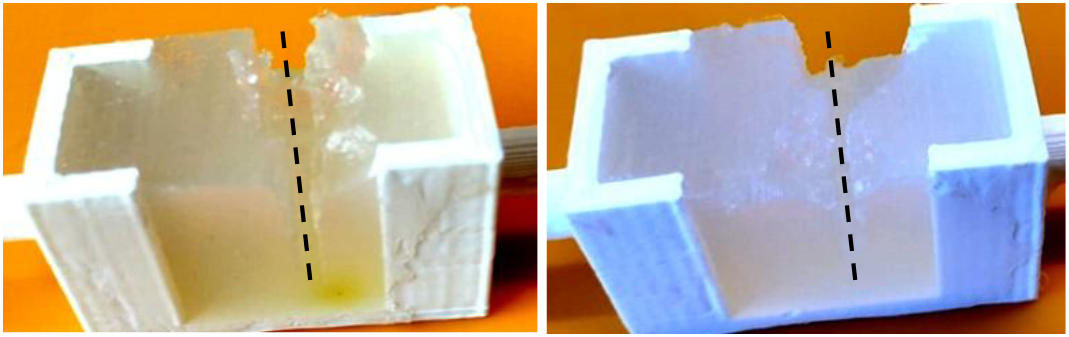
On the left, 3% agar after testing with BS blade. On the right, 3% agar after testing with B blade. Contrast adjusted. Dashed lines: blade path.

**Figure 17.**
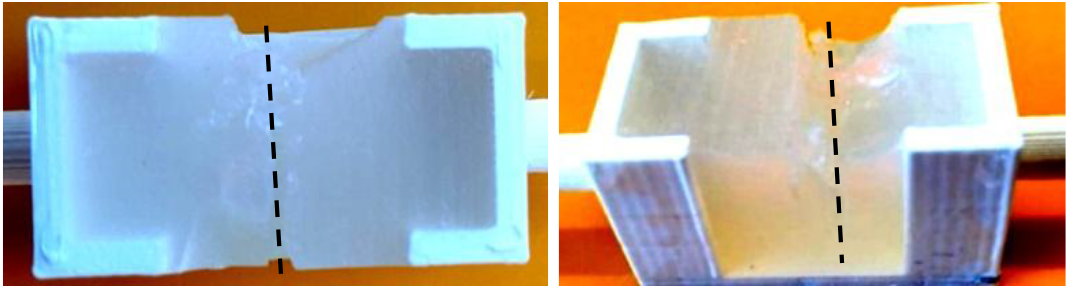
On the left, 3% agar after testing with S blade. On the right, 3% agar after testing with W blade. Contrast adjusted. Dashed lines: blade path.

With 5% agar there was a clear difference in the performance of the different blades. The serrulae of BS blades caused damage on the basal side of the substrate while the bump bit into the agar, ripping out material on the apical side of the substrate, as shown in Figure 18. The B and W blades caused minimal damage; only a small mark can be seen on the apical side of the substrate, as shown in Figure 19 and Figure 2.1 Some damage was caused by the serrulae of the S blades, as shown in Figure 20.

**Figure 18.**
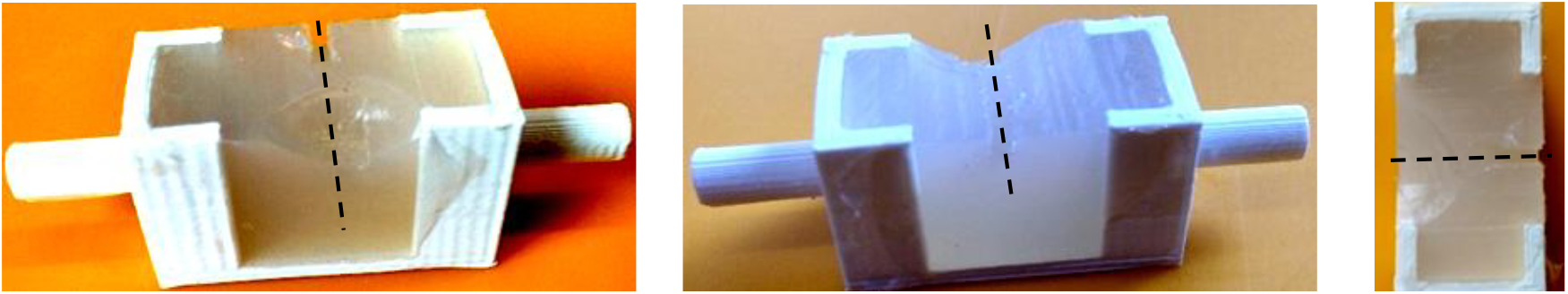
Different perspectives of 5% agar after testing with BS blade. Contrast adjusted. Dashed lines: blade path.

**Figure 19.**
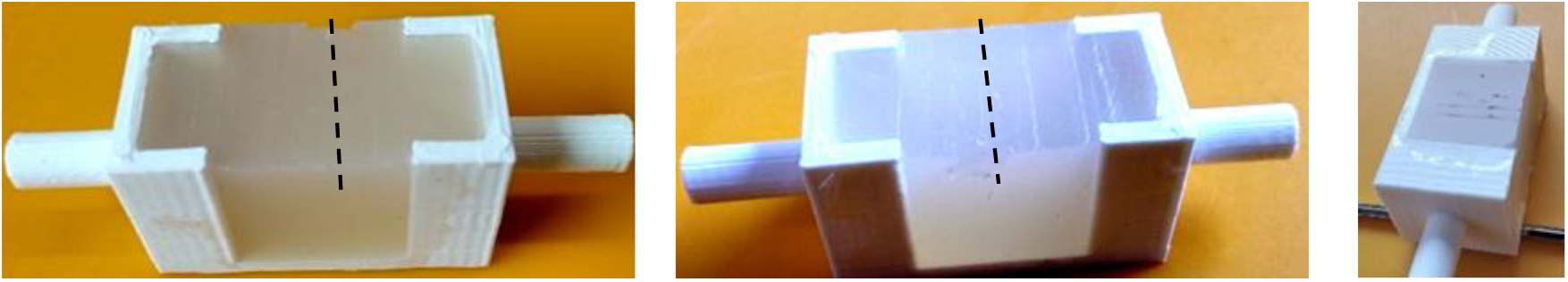
Different perspectives of 5% agar after testing with B blade. Contrast adjusted. Dashed lines: blade path.

**Figure 20.**
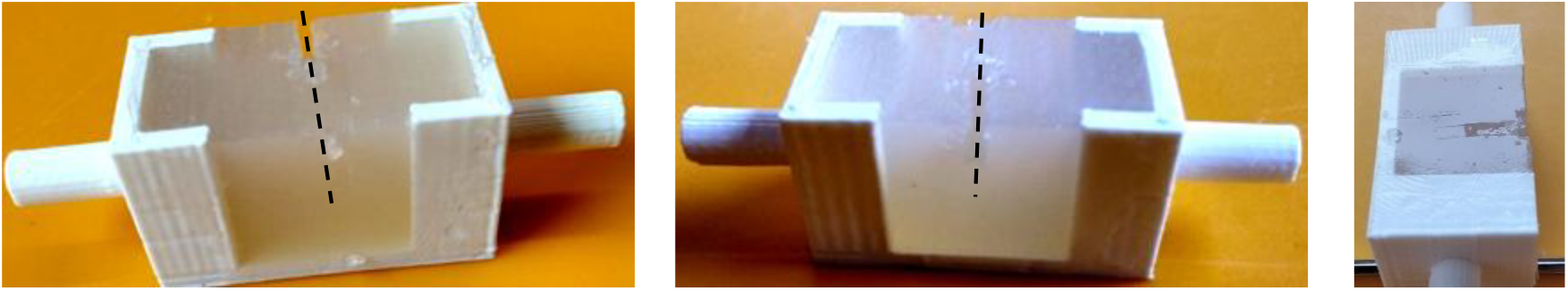
Different perspectives of 5% agar after testing with S blade. Contrast adjusted. Dashed lines: blade path.

**Figure 21.**
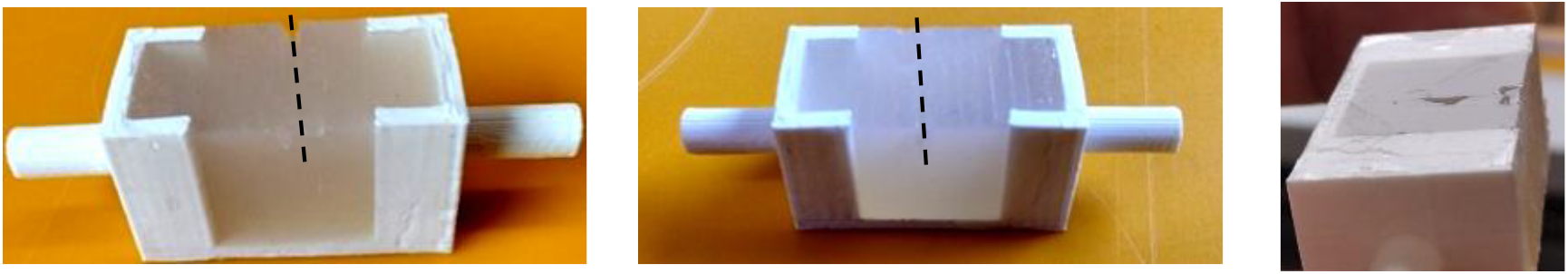
Different perspectives of 5% agar after testing with W blade. Contrast adjusted. Dashed lines: blade path.

Testing with 10% ballistic gelatine yielded very similar results to 5% agar. Only BS and S blades produced slight damage to the 10% agar, attributed to the serrulae, as shown in Figure 22. The W and B blades had negligible effect.

**Figure 22.**
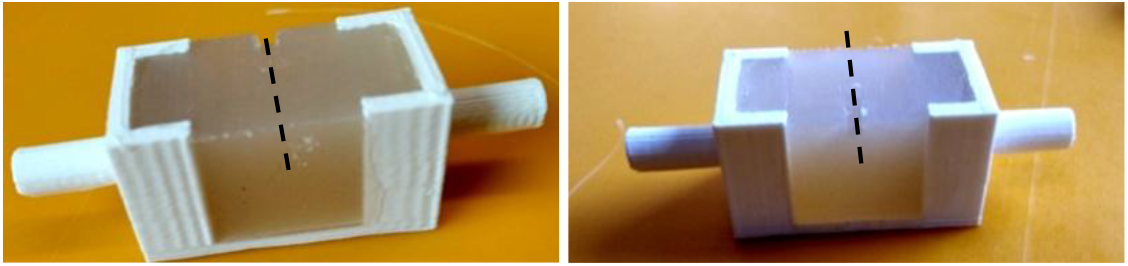
Different perspectives of 10% agar after testing with BS blade. Contrast adjusted. Dashed lines: blade path.

Using the “2D” test set up, pictures and recordings of the BS blades cutting into 5% agar substrates were obtained as shown in Figure 23.

**Figure 23.**
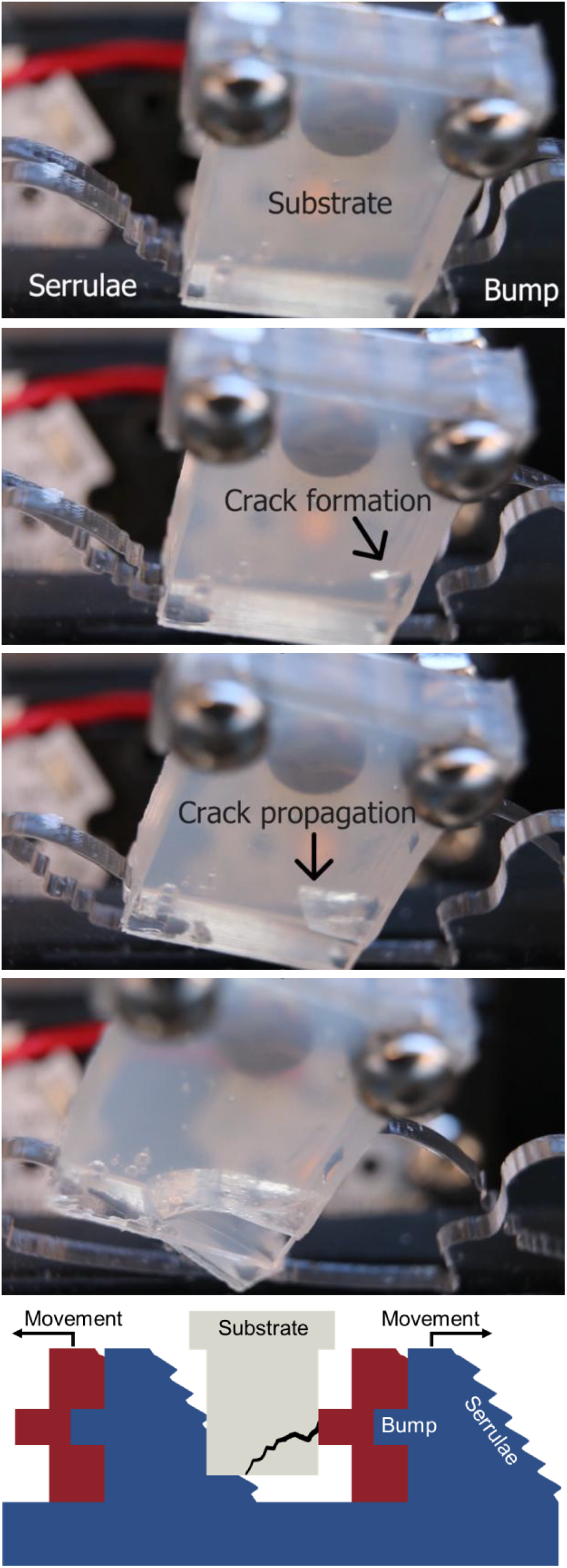
Top: Progression of substrate cutting showing crack formation and propagation during interaction with the bump and serrulae. Bottom: Schematic representation of the blade geometry and its interaction with the substrate.

When the “closed” set up was used, all the blades cut through all the substrates since the upper frame prevented substrate displacement out of the range of the teeth.

### 4.3 Added weight and change of scale cutting tests

Adding weight to the substrate carrier allowed the BS blades to bite into the 10% agar substrate and rip part of it off. The same happened when the scale was reduced to 200 as the bump could bite into the substrate (Table 2).

### 4.4 Influence of saw of another species

The cutting tests performed by the blades produced using the profile of the *H. brevis* ovipositor yielded results akin to those obtained for *R. scalaris* (Table 2).

### 5.1 Cutting test, added weight and change of scale

The cutting tests demonstrate that the presence of bump and serrulae increases the amount of damage on a substrate. In the first experiments, the BS blades were the only ones which could bite into the 5% agar as well as 10% ballistic gelatine while all blades could cut the 3% agar. A synergic effect between the bump and serrulae was demonstrated, since neither the S nor the B blades alone could damage the substrate to the extent caused by the BS blades. Whenever the blades were not able to cut the substrates, they were pushed out of the range of the teeth by the cutting motion, without suffering much damage.

Adding weight to the substrate carrier increased the vertical force of the blades against the substrate and allowed the BS blades to cut into the 10% agar, which is stronger. There is therefore a relationship between the ultimate stress of the substrate and the force against the substrate.

Reducing the scale of the teeth on the BS blades allowed the 10% agar substrate to be cut, which shows that the smaller the scale, the higher the ultimate strength of the substrate which can be cut.

Since when using the “closed” setting of the rig all the blades cut all the substrates in the same way, the only result was that provided there was enough force against the substrate and enough cut force, the system will cut all the substrates experimented with.

### 5.2 A sub-optimal mechanism?

There is no dual-bladed reciprocating cutting tool which presents a tooth geometry similar to that studied in this project. Intuitively, the optimal way to cause damage to the substrate using a reciprocating saw mechanism would be with teeth having the same bump characteristics on both sides (Figure 24, left). Those teeth would trap the substrate and damage it from both sides, without offering an inclined plane for the substrate to escape the cut. Such blades were indeed invented and patented for a reciprocating saw (Figure 24, right) [10].

**Figure 24.**
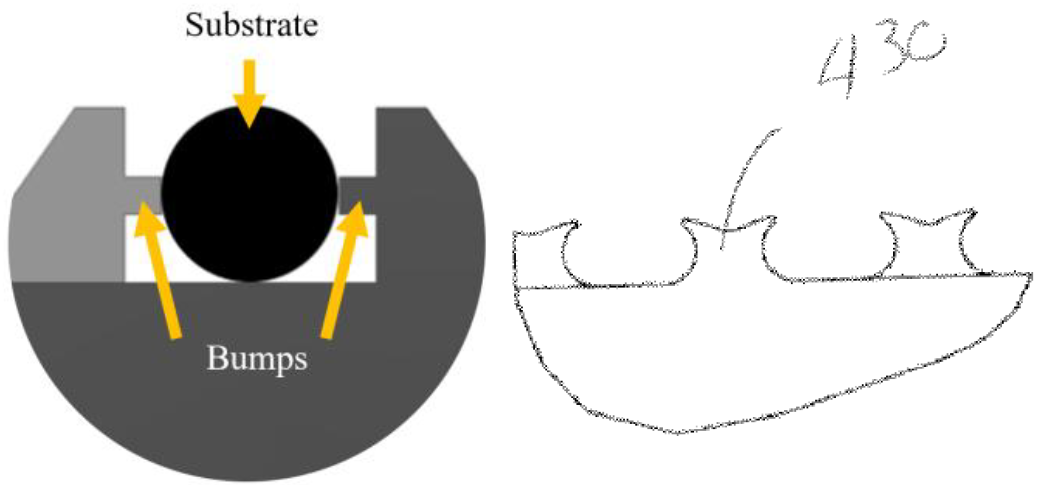
Left: Schematics of blades presenting bumps on both sides of the substrate. In light grey, the back blade; in dark grey the front blade. Right: Patented human design of teeth for a reciprocating saw. Reproduced from [10].

The question naturally arises as to whether this configuration is sub-optimal, in that it maximizes the damage done to the substrate. There should be a reason for this system to seem sub-optimal. Experimental results demonstrated that the different sets of blades, even the BS blades, did not cut all types of substrate. While it is clear that the bump and serrulae work together to damage a substrate of higher mechanical strength than would be possible with only one of the characteristics, let alone none, it is also clear that, above a certain threshold of substrate strength, the substrate cannot be damaged. Therefore, it would be reasonable to propose that the geometry of the studied teeth presents a threshold based on substrate strength above which the substrate is ejected.

### 5.3 Interaction between sawfly and plant

The sawfly cuts into the plant in order to lay eggs inside it, so that the eggs are protected and newly hatched larvae can feed on tissue close to where they hatch. This strategy requires that the surrounding plant tissue remains alive until the eggs hatch. Therefore, it is not in the sawfly’s interest to cause major damage to its host.

Most plants possess internal structures, such as vascular bundles. Damaging these would cripple the ability of the plant to survive and therefore, the sawfly needs to avoid damaging them while still being able to cut through the surrounding tissues. Since the vascular bundles are protected by strong fibres and highly lignified [11], a tooth is desirable that can cut the softer tissue but push the stronger tissue out of range. Furthermore, the sawfly must cut blindly, as the ovipositor is located ventrally close to the apex of the abdomen and the cutting is done mostly within the plant. The passive property of the selective cutting mechanism is therefore particularly helpful. The mechanism also seems to prevent damage to the ovipositor when it interacts with a strong substrate. In such a situation, this substrate would simply slip out of range, without much effect on the teeth, allowing the ovipositor to remain functional.

Comparing the shapes of teeth along the ovipositor of *R. scalaris*, the teeth at the base are adapted to cut the external layers of the plant (Figure 25 B). They possess the equivalent of a bump on each side of the tooth. This tooth geometry is closer to the human-made one mentioned above and is capable of causing more damage. The outer layer of the plant (epidermis) is stronger than the tissues beneath. Cutting the epidermis will not kill the plant, so the sawfly can use more aggressive teeth. In the central area of the ovipositor, (Figure 25 A), the teeth are adapted to cut selectively within the plant where the vascular bundles are located. In Figure 25 the ovipositor of *R. scalaris* (C) and a cross-section of stem of meadow buttercup, *Ranunculus acris* L. (D), which is one of the host plants of *R. scalaris*, are placed side by side and at the same scale. The zones presenting the two different kinds of teeth are shown, related to the kind of tissue they have adapted to cut or preserve (A and B).

**Figure 25.**
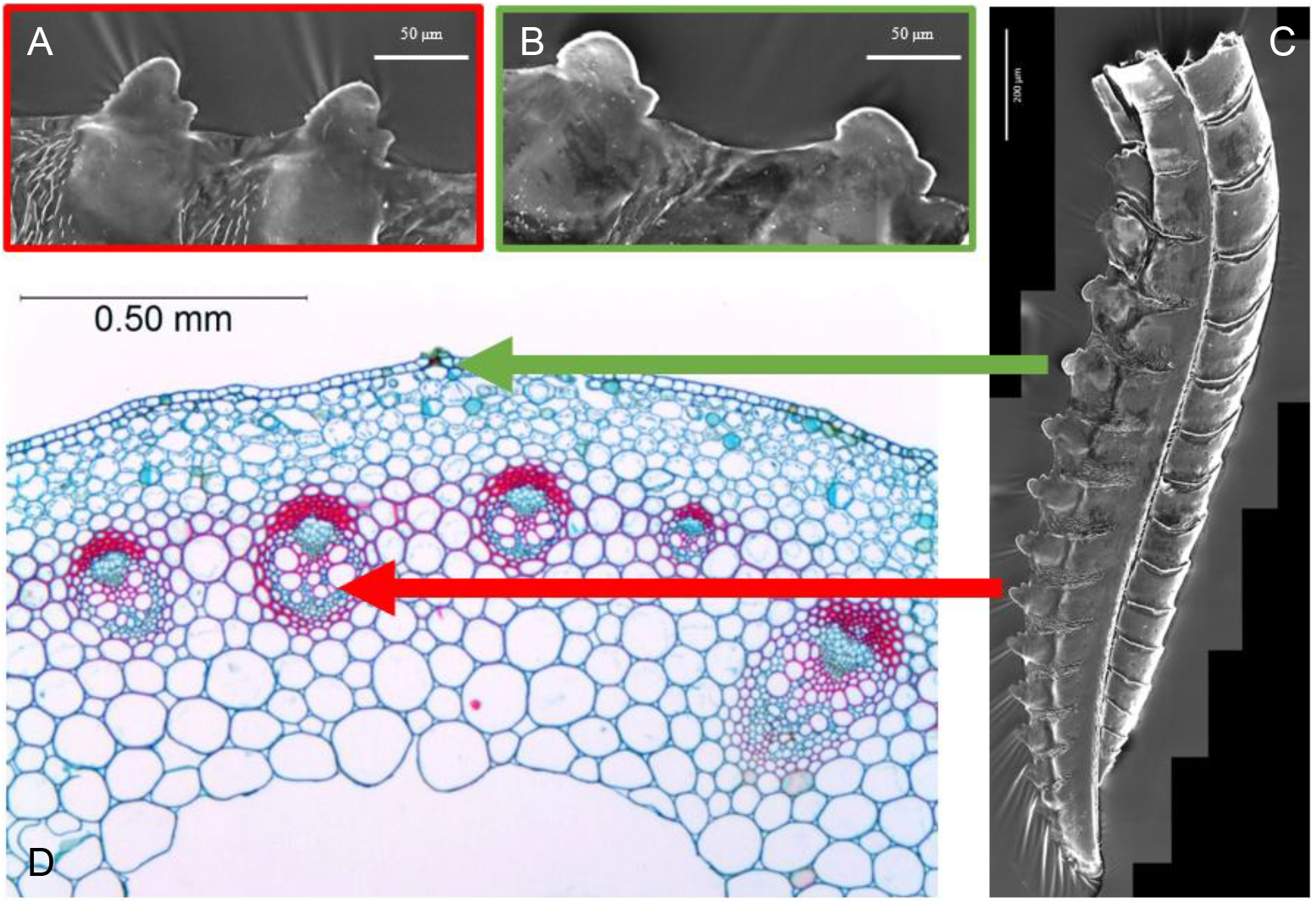
Relationship between teeth geometry and plant tissue. D: Cross-section of Ranunculus acris, adapted with permission from the author [12]. The areas presenting red cells are the vascular bundles. C: SEM image of R. scalaris ovipositor. B: Teeth in the basal region of the ovipositor not capable of selective cutting, more destructive. The green arrow connects the more destructive teeth to the outer layers of the plant. A: Teeth in the central region of the ovipositor capable of selective cutting. The red arrow connects the teeth capable of selective cutting to the area where vascular bundles are present.

The morphology of other ovipositors suggests that they share the same functional system. The ovipositor of *Pachyprotasis rapae* (Linnaeus, 1767), for example, also targets herbaceous plants and presents a similar bump-plus-serrulae structure. On the other hand, *Janus cynosbati* (Linnaeus, 1758), lays its eggs in oak shoots, which are much harder to kill, and its ovipositor has a very aggressive “double bump” tooth profile (Figure 26).

**Figure 26.**
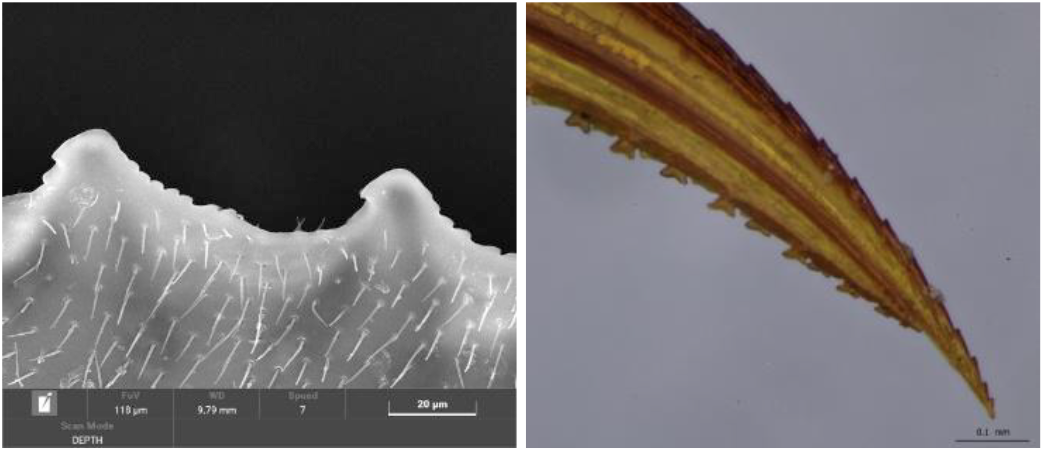
On the left lancet of Pachyprotasis rapae. On the right, lance and lancets of Janus cynosbati

Further tests were done using blades produced based on a tooth of *Hoplocampa brevis*. The behaviour of those blades in the tests was comparable to that in the tests done for *R. scalaris. H. brevis* lays its eggs in the calix of pear flowers, where the eggs hatch a couple of weeks later. The larvae eat the inside of the developing fruit [13]. This proves that the principle involved is not limited to a single species nor to exact morphology but works across species and with morphologies adapted to the species of plant being cut into.

### 5.4 Sawfly ovipositors as a model for surgical tools

We interviewed one surgeon and anonymously surveyed a further 14 surgeons who volunteered to participate. An extract of the interview follows. Each surgical operation requires a set of tools and, within this set, tools specific to that particular surgical procedure. Personal preference decides between scissors and electrosurgical scalpels. Electrosurgical scalpels, unlike scissors, cause coagulation on cutting, thus preventing bleeding if a blood vessel is sectioned accidentally. Scissors, unlike electrosurgical scalpels, do not damage the surrounding tissue when cutting. The general approach is to avoid cutting, but to move tissues aside. Ideally, once the epithelial tissue has been penetrated, only fat and connective tissue are cut until the target area is reached and the desired tissue is then cut.

When the working area is flooded with blood, the surgeons often struggle to see where they are working, where they are trying to get to, and where the blood is coming from. As a result, the chance of causing random damage increases. It is hard to stop the bleeding because the source is hard to find, and the risks increase as the duration of surgery lengthens. These situations are generally resolved by using electrosurgical scalpels to coagulate liquids while cutting, or suction tubes to aspirate the blood. The interviewed surgeon hinted that product design is more important than expected. This surgeon would love to experience the combination of the feeling of safety associated with electrosurgical scalpels with the elegance and accuracy of cutting possessed by scissors.

Fourteen surgeons, with an average of 10.7 years of experience, took part in the open-answer survey, responding to questions based on what was learned from the initial interview. 86% of the surgeons agree that the accumulation of blood impairs visibility of the targeted tissue and makes mistakes more likely. 57% prefer the use of electrocauterizing cutting tools to prevent bleeding. 79% express concerns regarding accidental tissue damage resulting from mistakes or thermal damage caused by electrocauterizing tools. Desire for a better product design was voiced by 57% of the respondents. The same percentage expressed concern about differentiating between organs and surrounding tissues, whether caused by impaired visibility, obesity or previous surgery.

In conclusion, both interview and survey highlighted a need for a safer tool, selective cutting of tissue and a desire for better product design, including flexibility, elegance, and user-friendliness. The passive selective cutting mechanism observed in sawfly ovipositors offers a promising avenue to inform the design of surgical tools that combine precision, safety, and selectivity.

## 6. Conclusions

The tests show the existence of a threshold which limits the substrates the blades can damage based on material strength. A connection between this ultimate stress threshold, the presence of bumps and serrulae, the force against the substrate, and the scale of the blade have been demonstrated. Studying the relationship between the ovipositors of sawflies and the plants in which they lay their eggs helps to understand the morphology of the ovipositor from a functional point of view. The tests performed indicate that the concept works across different species, which have adapted the teeth of their ovipositors to cope with a particular type of plant tissue.

This work identifies a bioinspired mechanism for passive, selective cutting, demonstrates its reproducibility across species, and highlights its potential value in tool design. Its translational potential is especially relevant to surgical applications, where precision and safety are critical.

## Acknowledgments

One of the authors, Dr Martí Verdaguer Mallorquí, acknowledges the financial support of the chool of Engineering & Physical Sciences at Heriot-Watt University through a doctoral scholarship. We thank the Senckenberg Deutsches Entomologisches Institut for providing the sawfly specimens used in this study.

We thank the surgeon who agreed to be interviewed and the 14 anonymous surgeons who participated in the questionnaire for their valuable insights into surgical tool use and design preferences. We thank Mihai Costea for granting permission to adapt the image of *Ranunculus acris* used in Figure 25.

## References

[1] Gao Y, Ellery A, Jaddou M, et al. Planetary micro-penetrator concept study with biomimetric drill and sampler design. IEEE Trans Aerosp Electron Syst. 2007;43(3):875– 885.

[2] Frasson L, Ko SY, Turner A, et al. STING: a soft-tissue intervention and neurosurgical guide to access deep brain lesions through curved trajectories. Proc Inst Mech Eng H. 2009;224(6):775–788.

[3] Verdaguer Mallorquí M, Vincent J, Liston A, et al. Knowledge from hymenopteran ovipositors: a review of past and current biomimetic research. Bioinspir Biomim. 2025;20(031001).

[4] Weltz CE, Vilhelmsen L. The saws of sawflies: exploring the morphology of the ovipositor in Tenthredinoidea (Insecta: Hymenoptera), with emphasis on Nematinae. J Nat Hist [Internet]. 2014;48(3–4):133–183.

[5] Goulet H. Nearctic Tenthredo: a monograph of the verticalis and prosopa groups (Hymenoptera: Tenthredinidae). Nova Supplementa Entomologica. 2020;26(2):1–178.

[6] Viitasaari M. The suborder Symphyta of the Hymenoptera. Sawflies. 2002;11–174.

[7] Maiden NR, Fisk W, Wachsberger C, et al. Ballistics ordnance gelatine - How different concentrations, temperatures and curing times affect calibration results. J Forensic Leg Med. 2015;34:145–150.

[8] Pullen A, Kieser DC, Hooper G. Ballistic gelatin calibration standardisation. BMJ Mil Health. 2022;168(2):124–127.

[9] Subhash G, Liu Q, Moore DF, et al. Concentration dependence of tensile behavior in agarose gel using digital image correlation. Exp Mech. 2011;51:255–262.

[10] Vitantonio M, Jackson T, Nottingham J, et al. Dual blade reciprocating saw. Australian patent office; 2012.

[11] Gibson LJ. The hierarchical structure and mechanics of plant materials. J R Soc Interface. 2012;9(76):2749–2766.

[12] M. Costea. Phyto Images: Ranunculus acris [Internet]. 2012. Available from: http://phytoimages.siu.edu/imgs/Cusman1/r/Ranunculaceae_Ranunculus_acris_45642.html.

[13] Liston A, Prous M, Vårdal H. A review of West Palaearctic Hoplocampa species, focussing on Sweden (Hymenoptera, Tenthredinidae). Zootaxa. 2019;4615(1):1–45.

